# *Legionella* becoming a mutualist: adaptive processes shaping the genome of symbiont in the louse *Polyplax serrata*

**DOI:** 10.1101/155820

**Authors:** Jana Říhová, Eva Nováková, Filip Husník, Václav Hypša

**Author notes:** Corresponding author: Václav Hypša.

## Abstract

Legionellaceae are intracellular bacteria known as important pathogens of man. In the environment, they are mainly found in biofilms associated with amoebas. In contrast to another gammaproteobacterial family Enterobacteriaceae which established a broad spectrum of symbioses with many insect taxa, the only instance of legionella-like symbiont has been reported from lice of the genus *Polyplax*. Here, we sequenced the complete genome of this symbiont and compared its main characteristics to other *Legionella* species and insect symbionts. Based on rigorous multigene phylogenetic analyses, we confirm this bacterium as a member of the genus *Legionella* and propose the name *Candidatus* Legionella polyplacis, sp.n. We show that the genome of *Ca.* Legionella polyplacis underwent massive degeneration, including considerable size reduction (529.746 bp, 484 protein coding genes) and a severe decrease in GC content (23%). We identify several possible constraints underlying the evolution of this bacterium. On one hand, *Ca.* Legionella polyplacis and the louse symbionts *Riesia* and *Puchtella* experienced convergent evolution, perhaps due to adaptation to similar hosts. On the other hand, some metabolic differences are likely to reflect different phylogenetic positions of the symbionts and hence availability of particular metabolic function in the ancestor. This is exemplified by different arrangements of thiamine metabolism in *Ca.* Legionella polyplacis and *Riesia*. Finally, horizontal gene transfer is shown to play a significant role in the adaptive and diversification process. Particularly, we show that *Ca.* L. polyplacis horizontally acquired a complete biotin operon (bioADCHFB) that likely assisted this bacterium when becoming an obligate mutualist.

## Introduction

Legionellaceae are mainly known as important bacterial pathogens of man (Diederen 2008), although their life strategy is generally bound to biofilms where they live in intracellular symbiotic associations with amoebas and other protists (Fields 1996). Consequently, all known species have therefore been described either from water sources or clinical materials. The only known exception to this rule is the obligate symbiont of the lice *Polyplax serrata* and *P. spinulosa* originally characterized by light and electron microscopy (Ries 1931; Volf 1991). Based on the 16S rDNA sequence, this bacterium was later on suggested to be a member of the genus *Legionella* (Hypsa and Krizek 2007). In this work, the transition from a typical legionella to an obligate symbiont was inferred from the presence of the bacterium in all tested louse individuals, suggesting their transovarial transmission, and from the typical shift in GC content. These traits are common for many obligatory symbionts living intracellularly in various insects. Typically, such bacteria reside in specialized organs, usually called bacteriomes, and are presumed to supply the host with some essential compounds, mainly vitamins and amino acids, missing in their diet (Douglas 1989). Their evolution/adaptation towards this role is usually accompanied by extensive modifications of their genomes, consisting mainly of gene (or complete function) losses and decrease of GC content (Woolfit and Bromham 2003). While the latter is considered a sign of relaxed selection, the reduction of metabolic capacities results mostly from adaptive processes. In the course of evolution, these bacteria lose metabolic functions fulfilled by the host and retain (sometimes even acquire by horizontal transfer; Husnik, et al. 2013; Nakabachi, et al. 2013; Nikoh, et al. 2014) functions essential for the host’s development and/or reproduction. Due to these processes, the bacteria can evolve an array of different life strategies, from parasites/pathogens to obligate symbionts. It was for example demonstrated that the notorious insect parasite *Wolbachia* turned into a mutualistic bacterium in the bedbug *Cimex lectularius* upon horizontal acquisition of the operon for biotin biosynthesis (Nikoh, et al. 2014; Gerth and Bleidorn 2017).

Several decades lasting research on insect-bacteria symbiosis revealed broad variety of these associations and resulted in many fundamental discoveries (Moran 1996; Ochman and Moran 2001; McCutcheon and Keeling 2014; Moran and Bennett 2014). One of the interesting outcomes is the broad diversity of bacteria which are generally capable to establish obligate symbiosis with insects, including such diverse taxa as Enterobacteriales, Rickettsiales, Blattabacteriaceae, Flavobacteriales, etc. However, within this broad bacterial diversity, several groups show particular tendency to this behavior (e.g. Enterobacteriaceae, more specifically the genera *Arsenophonus* and *Sodalis*), while others are found less frequently. Legionellaceae is one of these rare groups, with the *Polyplax*associated bacterium being the only case of such intracellular obligate insect symbiont. This raises an interesting question on how this bacterium established its unique symbiotic relationships with lice and what changes it underwent during its adaption to obligate symbiosis. In this respect, it is important that there is a solid body of genomic information for both the genus *Legionella* and many other bacterial symbionts from various blood sucking insects. In a recent genomic work, comparing 38 *Legionella* species, Burstein, et al. (2016) revealed surprising diversity of their genomic traits, including major differences in the most general characteristics such as genome size or GC content. The main focus of this genomic comparison, particularly important in respect to the evolution of legionellae virulence, were effectors, especially those responsible for entering the host cell. The computational analysis inferred several thousands of candidate effectors and indicated that many of these factors have been recently acquired by horizontal transfer. This shows the genus *Legionella* as highly dynamic system with ongoing genome rearrangements and adaptations towards the intracellular lifestyle. The newly characterized legionella-like louse symbiont thus provides us with a unique opportunity to study the processes of the genomic changes and compare them to “free living” *Legionella* on one hand and to the unrelated enterobacterial symbionts on the other hand. We demonstrate that the symbiotic bacterium originated within the genus *Legionella*. We also show that its genome followed an evolutionary route typical for obligatory symbionts, and underwent a surprisingly convergent evolution with unrelated enterobacterial symbiont from the hominid lice. Finally, we show that its adaptation to the role of obligatory symbiont included horizontal acquisition of six genes coding a biotin operon, in analogy to the mutualistic *Wolbachia* from bedbugs.

## Materials and Methods

### Sample preparation

The specimens of *Polyplax serrata* were collected during autumn 2011 from *Apodemus flavicolis* mice trapped around Baiersbronn, Germany and stored in absolute ethanol at -4^o^C. Since the obligate bacterial symbionts are uncultivable outside the host cells, total DNA was obtained by extraction from whole abdomens of 25 louse individuals (QiaAmp DNA Micro Kit, Qiaqen). DNA concentration was assessed by the Qubit High Sensitivity Kit (Invitrogen) and 1% agarose gel electrophoresis.

### Genome sequencing and assembly

The *Polyplax* sample was sequenced on one lane of Illumina HiSeq2000 (GeneCore, Heidelberg) using 2 × 100 paired-end reads (PE) library with an insert size of 150 bp. After checking and filtering for the quality, the resulting data set contained 309,892,186 reads in total. The reads were assembled using the SPAdes assembler v 3.10 (Bankevich et al. 2012), with the parameter *careful*, decreasing number of mismatches and indels. To check for possible presence of bacterial plasmid(s) in the data, we submitted complete assembly to the PlasmidFinder (Carattoli et al. 2014) with sensitivity set to three different thresholds (95%, 85%, and 60%). Phylogenetic affiliations of the contigs were determined by PhylaAMPHORA (Wu and Scott 2012). Of the total 124,985 contigs, 112 were assigned to the order Legionellales. Trimmed reads were mapped on these contigs and filtered using BWA v. 0.7.15 (Li and Durbin 2009) and retrieved by Samtools (Li, et al. 2009). This set of reads was subsequently assembled by two alternative assemblers, SPAdes and A5 assembly pipeline (Coil, Jospin and Darling 2015). The latter software produced a closed 529,746 bp long genome. Its quality was checked and base calls were polished by Pilon v1.20 (Walker et al. 2014).

### Genome annotation

The genome was annotated using two tools services: RAST (Aziz, et al. 2008), and PROKKA (Seemann 2014). The complete genome was deposited in GenBank with the accession number SAMN07171993. To scan for potential horizontal gene transfer(s) (HGT), we retrieved 50 most similar sequences for each protein using the Blastp algorithm (Altschul et al. 1990) against the nr (non-redundant) protein database. Metabolic pathways for B vitamins were reconstructed using KEGG Mapper (http://www.genome.jp/kegg/mapper.html). The absence of genes in important metabolic pathways was verified using Blast searches.

### Phylogenetic analyses

Sixty four proteins for a multigene analysis were selected based on the recent genomic study by (Burstein et al. 2016) and their orthologs in the *L. polyplacis* were determined using Blastp search against the *L. polyplacis* genome. The sequences were aligned using MAFFT v. 1.3.5 (Katoh et al. 2002) implemented in the software Geneious (Katoh and Standley 2013). Equivocally aligned positions and divergent regions were eliminated by GBlocks (Castresana 2000). The alignments were created separately for each protein and multigene matrix was obtained by their concatenation. To minimize phylogenetic artifacts caused by rapid evolution and nucleotide bias of the symbiont sequences, we used PhyloBayes MPI v. 1.5a (Lartillot et al. 2013) with the CAT-GTR model and dayhoff66 amino acid recoding, and run it for 32,000 generations. This approach has been previously shown to decrease phylogenetic artifacts affecting branching of symbiotic bacteria in bacterial phylogenies (Husnik et al. 2011).

For the candidate HGTs, i.e. the biotin operon genes (see Results and Discussion), we prepared a representative set of orthologs covering several bacterial groups and used the alignment method described above to build amino acid matrices. The best evolutionary model for all matrices, determined in Prottest 3.2 (Darriba et al. 2011) by Akaike information criterion (AIC), was LG with a proportion of invariable sites and evolutionary rates separated in four categories of gamma distribution (LG+I+G). Phylogenetic reconstructions were done by maximum-likelihood analyses with 100 bootstrap replicates using PhyML v. 2.2.0 (Guindon et al. 2010) for each gene separately and also for a concatenated matrix of all six genes. Posterior probabilities for individual branches were determined by MrBayes v 3.2.6 (Huelsenbeck and Ronquist 2001) with the same evolutionary model (LG+I+G) and remaining parameters determined by the analysis. The analysis was run in 4 default chains for 10,000,000 generations. Convergence of all Bayesian analyses was checked in Tracer v1.6.0 (Rambaut et al. 2014), and for MrBayes also by the values of standard deviation of split (below 0.01) and PSRF+ (reached the value 1.0). Based on the availability of the analyzed gene, one of the following bacteria was used as an outgroup: *Kurthia* sp. (Firmicutes), *Geobacter sulfurreducens* (Deltaproteobacteria) and *Cyanothece* sp. (Cyanobacteria). Graphical representation of the trees was processed in FigTree (http://tree.bio.ed.ac.uk/software/figtree/) and Inkscape (http://www.inkscape.org/).

### Comparative genome analyses

In order to reveal general similarity patterns and possible functional convergences between the symbiont of *P. serrata* and other symbiotic bacteria, we have treated the genomes and genes as communities and their components, and employed NMDS clustering analysis routinely used in microbial ecology. The dissimilarity matrix based on Bray-Curtis distances was calculated for 4632 GOCs from 94 bacterial genomes (Supplementary file 1 Table S2). Draft genomes of poor quality have not been included. In particular we have omitted two genomes of high interest, i.e. *Riesia pediculischaeffi* and *Sodalis* endosymbiont of *Proechinophthirus fluctus* (Boyd et al. 2016). All the genome data were retrieved as proteomes from GenBank and assigned to particular functional orthologs as defined by the COG database (Tatusov et al. 2000) using rpsblast (Altschul et al. 1990). While the dataset includes free living Legionellaceae with large genomes, the matrix was transformed using a percentage proportion for each COG ortholog calculated from the COG sum of each genome. The matrix of 4632 orthologs was converted into the biom format. The Bray Curtis dissimilarities were calculated and the NMDS analysis was performed using vegan package functions in R (Dixon 2003). In order to show the genome functional similarities among lice symbionts with distinct phylogenetic origin, a Neighbour Joining (NJ) tree was calculated in T-rex (Alix et al., 2012) using the same distance matrix.

### Proposal for the species name of Legionella-like endosymbiont

As we show that the symbiont clusters within the genus *Legionella*, we propose in accordance with the terms for species designation which have not been cultivated in a laboratory media the name “*Candidatus* Legionela polyplacis” sp. nov. (hereafter *Legionella polyplacis* for simplicity). The specific name “polyplacis” refers to the genus of its insect host, the louse *Polyplax serrata*. The bacterium is the only member of the genus for which endosymbiotic association with insects has been documented.

## Results and Discussion

### Genome properties

The complete genome of *Legionella polyplacis* is 529 746 bp long, has extremely low CG content (23%) and coding density 84.8%. According to the RAST annotation (Supplementary file 1 Table S3), it contains 484 protein encoding genes (pegs) with the average length of 939 bp, 3 genes coding for rRNA and 36 genes for tRNAs. In two cases, pairs of adjacent pegs, annotated by identical names, corresponded clearly to a gene interrupted by a stop codon (DNA polymerase I (EC 2.7.7.7) and COG1565: Uncharacterized conserved protein). Since we confirmed the presence of the stop codons by mapping the reads on the assembled genome, we suppose that these cases represent either a functional split into two separate genes or an early stage of pseudogenization. To avoid an arbitrary determination of other possible pseudogenes, based on the sequence lengths, we provide in the Supplementary file 1 Table S3 the comparison of length ratios between the *L. polyplacis* genes and their closest Blast hits. In summary, of the 484 coding genes, only 48 are shorter than 90% of the blast-identified closest homologs, 23 of them shorter than 80%. The genome size and gene number place *L. polyplacis* among many other obligate symbionts of insects. In Table 1., we show the comparison of these general genomic characteristics between typical legionellae, *L. polyplacis* and other hominid louse symbionts.

Of the coding genes, 105 contain transmembrane helices and 4 signal peptides. No CRIPSR repeats were identified. PlasmidFinder, run under the sensitivity range 95% - 60%, did not reveal any candidate plasmid sequence. Majority of the coding genes (445 genes) could be assigned by Blastp unequivocally to the genus *Legionella* (i.e. 5 best hits corresponded to the genus *Legionella*), for 20 genes the search returned *Legionella* together with some other bacterial genera, for 30 genes we either did not get any match or the hits corresponded to other bacteria than *Legionella*. Among the “*non-legionella”* genes, some were highly conservative (ribosomal proteins) or very short genes difficult to assign based on the blast algorithm. However, six of these “*non-legionella*” genes formed a complete biotin operon known from several nonrelated symbiotic bacteria, suggesting its horizontal acquisition in the *L. polyplacis*). A significance of this HGT in respect to the symbiotic nature of this legionella is discussed below. The genes with no blast hits were mostly annotated as hypothetical proteins, usually with very short sequences. Of the metabolic functions considered particularly significant in the endosymbiotic bacteria, we detected several complete pathways (Fig 1) and the horizontally acquired capability of biotin synthesis (Fig 2).

**Figure 1.**
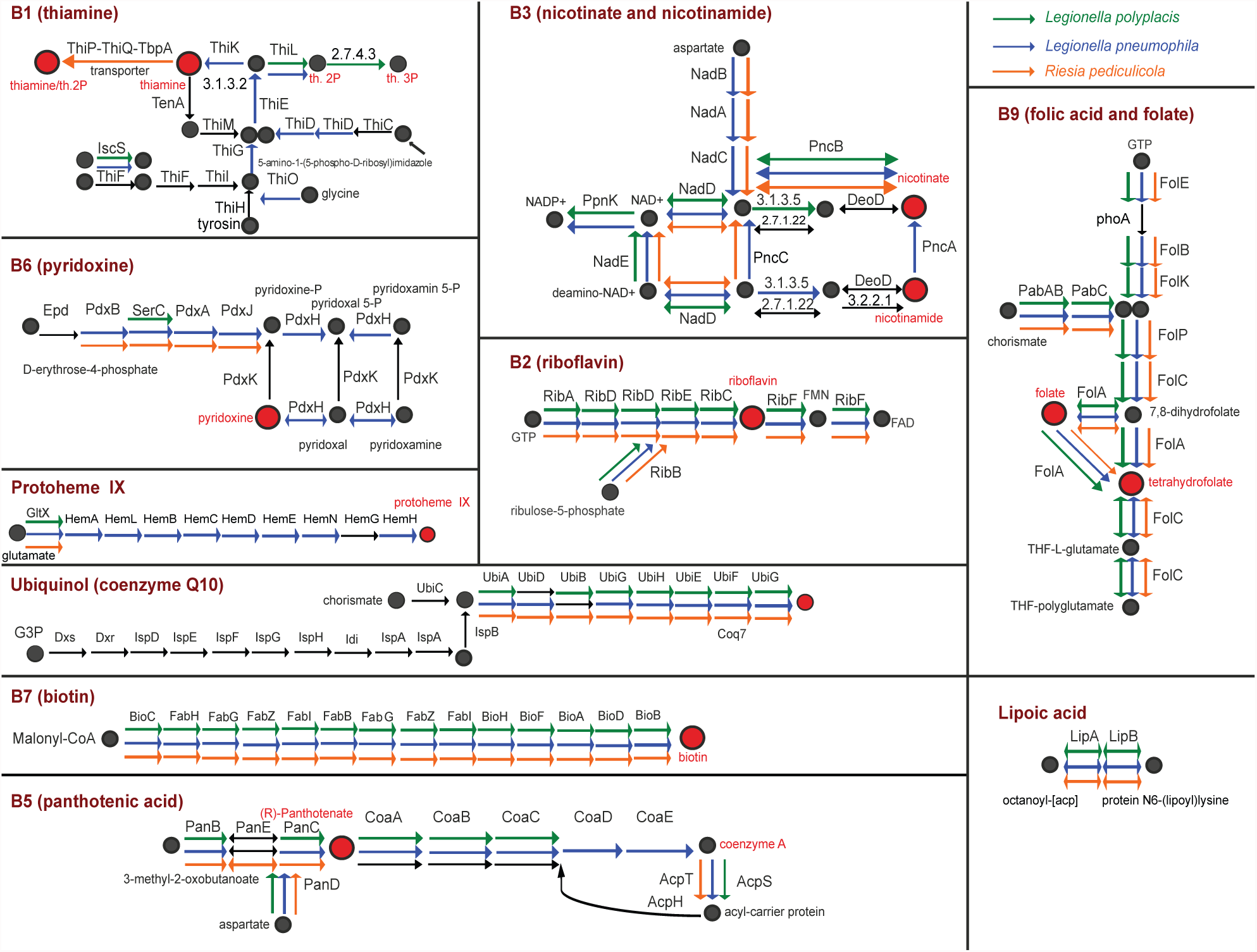
Comparison of B-vitamins and cofactor pathways for *Ca*. Legionella polyplacis, *Riesia pediculicola* and *Legionella pneumophila*. Black arrows designate missing genes.

**Figure 2.**
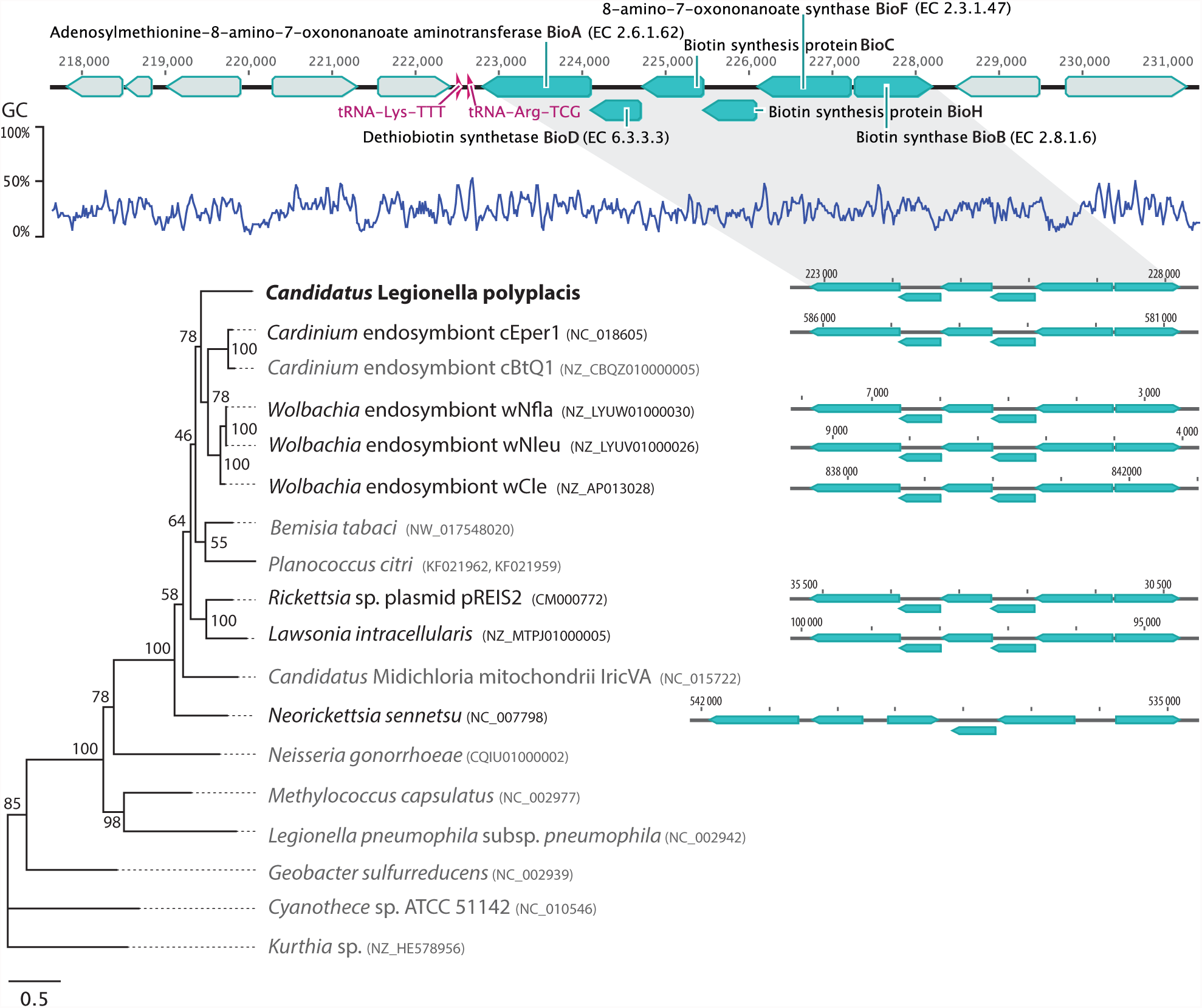
Structure of the horizontally acquired biotin operon in *Ca*. Legionella polyplacis and its putative evolutionary origin, based on ML analyses of all available genes. Blue arrowhead blocks represent genes for biotin synthesis organized in an intact operon. The adjacent genes are as follows: for BioA from right to left: hypothetical protein, Isopentenyl-diphosphate delta-isomerase FMNdependent, 4-hydroxy-tetrahydrodipicolinate synthase, 4Fe-4S ferredoxin, TsaB protein; for BioB from left: Aminomethyltransferase (glycine cleavage system T protein), and Glycine dehydrogenase [decarboxylating] (glycine cleavage system P2 protein). The position of the complete operon is visualized in the respective genomes (black font). The species with missing genes or disrupted operon structure (*Legionella pneumophila*) are in gray. The numbers at the tree nodes stand for the bootstrap values.

### Phylogeny and overall genomic evolution

The multigene analyses confirm that *L. polyplacis*, the symbiotic bacterium from the lice of the genus *Polyplax,* have indeed originated within the genus *Legionella* (Fig 3, Supplementary file 2), as previously suggested based on 16S rDNA (Hypša and Křížek, 2007). Although the Phylobayes analysis did not converge even after 32,000 generation, it yielded an identical result with the Phyml analysis in respect to the *L. polyplacis* position. In both analyses, it clustered on an extremely long branch, but with the strong support, within the group corresponding to the *L. micdadei* clade of Burstein et al. (2015). This position was further supported by the Blast analysis of the genes conserved in all species, including the symbiont, where the first eight species belonged to the *L. micdadei* clade ( i.e. *L. feeleii, L. lansingensis, L. brunensis, L. hackeliae, L. jamestowniensis, L. maceachernii, L. jordanis* and *L. nautarum*). The basic structure of the whole *Legionella* cluster (i.e. monophyly of the main groups) corresponds to that reported by Burstein et al. (2015). The Phyml-derived topology retains closer similarity to their ML based tree. This general agreement among results of our ML, Bayesian inference based on the recoded matrix, and the Burstein’s et al. (2015) topologies indicate that the data provide a reliable information for reconstructing the relationships within this group. Comparing the genome size of *L. polyplacis* (529 746 bp and 484 coding genes) to that determined for other legionellae shows that similarly to other insect symbionts, the genome of *L. polyplacis* experienced considerable reduction in course of its adaptation to the symbiotic lifestyle. Although the 38 *Legionella* spp. compared by Burstein et al., (2016) varied considerably in their genome sizes (2MB-5MB, 2,000-4500 genes), they never approached the degree of reduction determined for the *L. polyplacis* genome. Due to this dramatic reduction, *L. polyplacis* lost many complete pathways and systems present in other *Legionella* spp.

**Figure 3.**
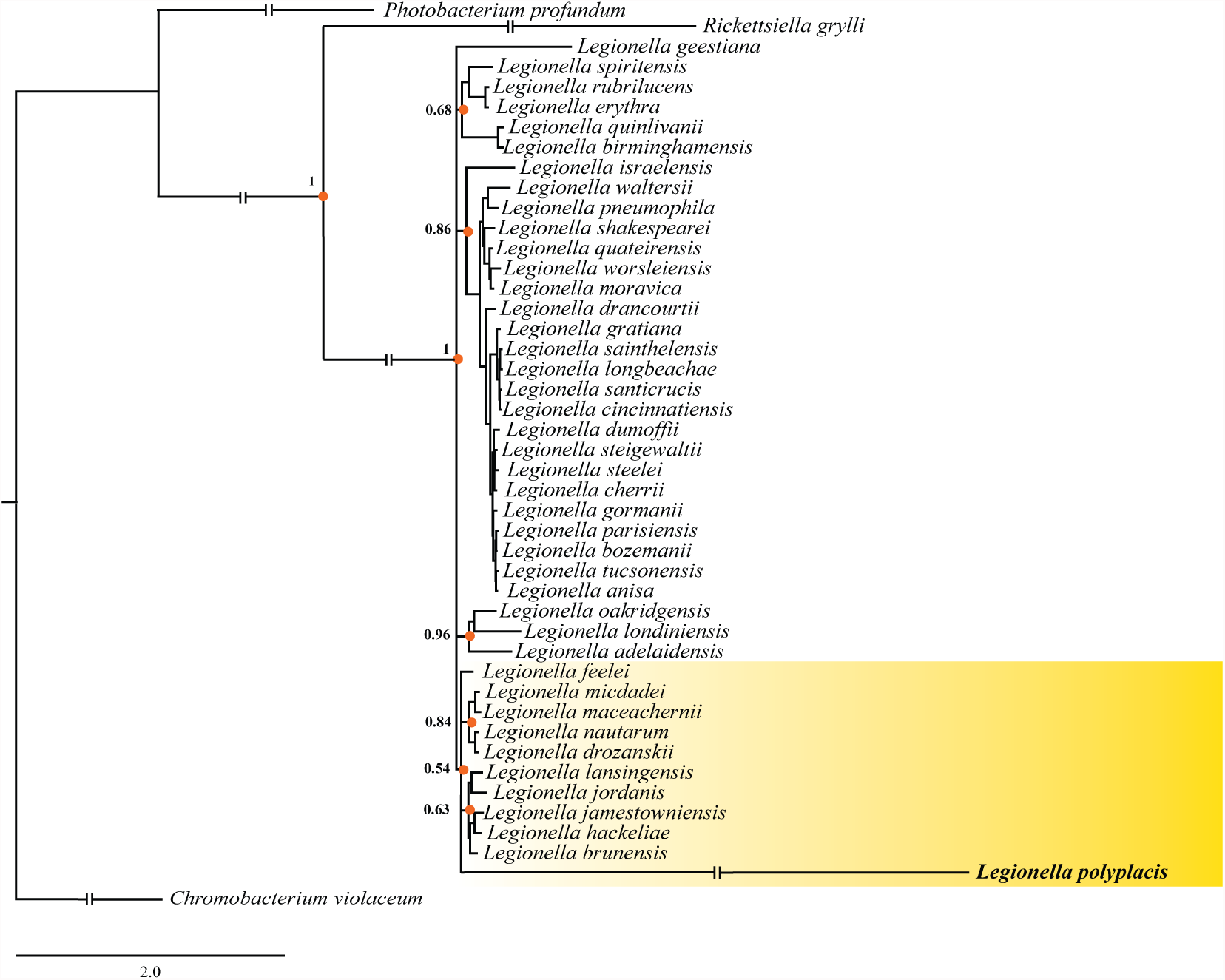
Phylogenetic tree inferred by PhyloBayes analysis of concatenated 64-gene matrix. The orange dots (with posterior probability values) highlight arbitrary selected monophyletic clusters as main components of the tree structure. The interrupted branches were shortened by 50%.

Since nothing is known about the origin of bacterial symbionts in *Polyplax* species other than *P. serrata* and *P. spinulosa*, it is difficult to estimate the time frame of this reduction. In phylogenetic study published by Light et al., (2010), the family Polyplacidae are revealed as a paraphyletic taxon, with the genus *Polyplax* branching as sister group to a Pediculidae/Phthiridae/Pedicinidae cluster. The estimated diversification time for these two groups, around 45 Mya, can thus be considered an upper limit for the origin of *L. polyplacis*. This would place *L. polyplacis* close to another obligatory symbiont associated with blood sucking insect, namely *Wigglesworthia glossinidia*, with the estimated origin of 40 Mya (Naito and Pawlowska 2016). However, since the *Polyplax*’s long branch in the Light et al. (2010) analysis indicates that other closer relatives of *Polyplax* might be missing in the taxa set, this time frame may be considerably overestimated. For example, another louse symbiont, *Riesia pediculicola* from the human louse, was shown to had reached similar genome state within approximately 13-25 Mya (Boyd, et al. 2014).

Regardless the uncertainty in time estimates, *L. polyplacis*, *R. pediculicola* and the other recently described symbionts of lice (Boyd, et al. 2017) reached similar basic genome characteristics (Table 1). More interestingly, the NMDS analysis (Fig 4) and particularly the distance based NJ tree (Fig 5) placed *L. polyplacis* genome far from other legionellae but remarkably close to the phylogenetically distant genus *Riesia* (i.e. member of the *Arsenophonus* cluster). In the NJ tree, all of the louse symbionts, including *Puchtella*, even form a monophyletic lineage. It is important to notice that this clustering is not determined by the genome size as the other highly modified symbionts are scattered across the whole NMDS plot. Even more interestingly, the primary symbionts associated with blood feeding pupiparans, i.e. *Wigglesworthia* spp., *Arsenophonus melophagi* and *A. lipopteni*, cluster within a distant independent group relatively close to each other. In other words, the symbionts of blood feeding insects show clear tendency towards clustering according to their host phylogeny (and hence perhaps biology) rather than their own phylogeny. Although the sample of the symbionts from the blood-feeding insects is small for any decisive inference, the pattern suggests that the clustering may reflect more subtle aspects than just general genome reduction and the host’s source of nutrients.

**Figure 4.**
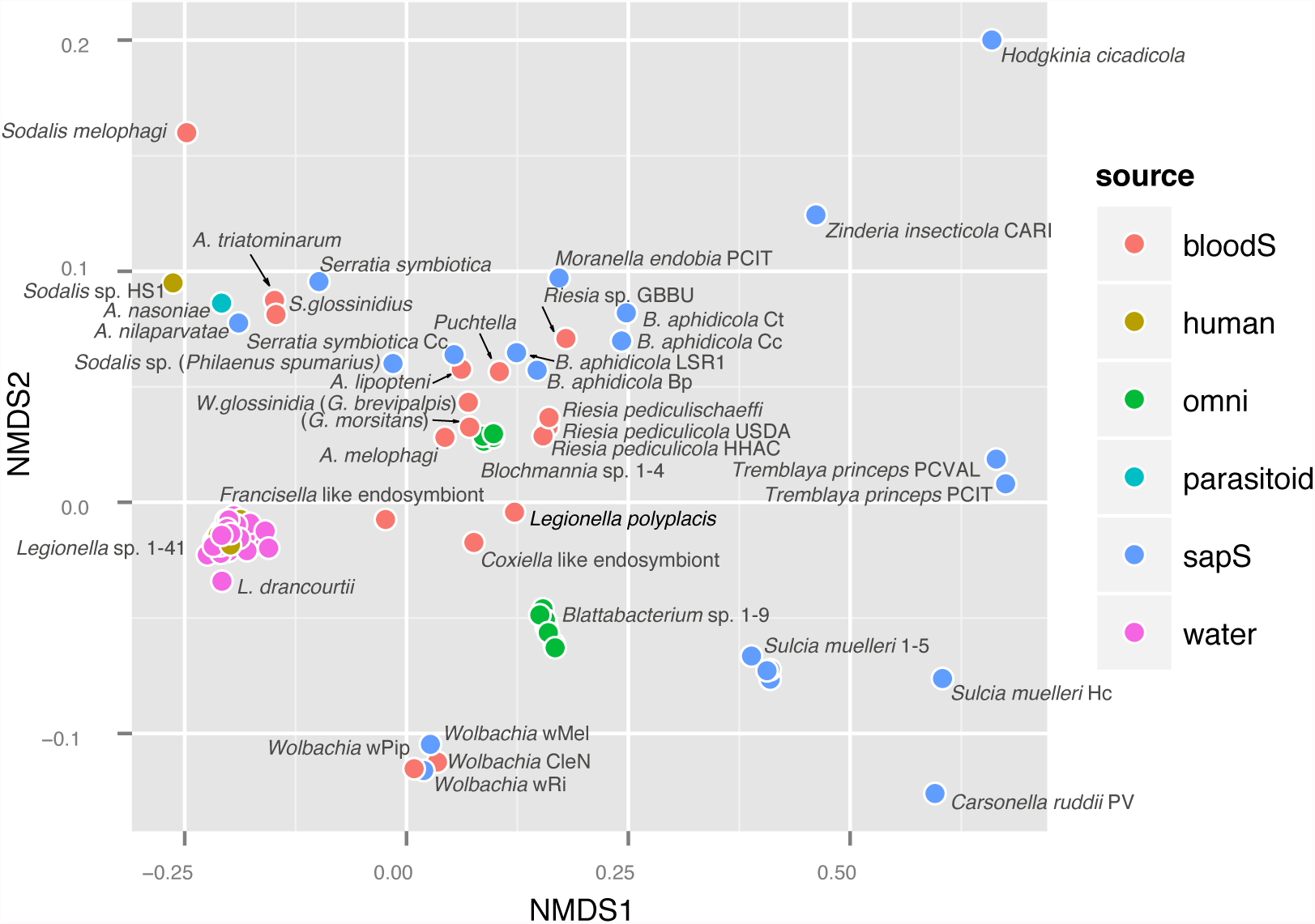
NMDS analysis based on Bray-Curtis dissimilarities calculated for the genome content of selected bacteria analyzed across all COG orthologs. The legend abbreviations for the bacterial source are as follows: **bloodS**: blood sucking host, **human**: clinical isolate, omni: omnivorous insect host, **parasitoid**: i.e. *Nasonia vitripennis* host, **sapS**: sap sucking insect host, **water:** environmental water sample. List of the analysed genomes is provided in Supplementary file 1 Table S2.

**Figure 5.**
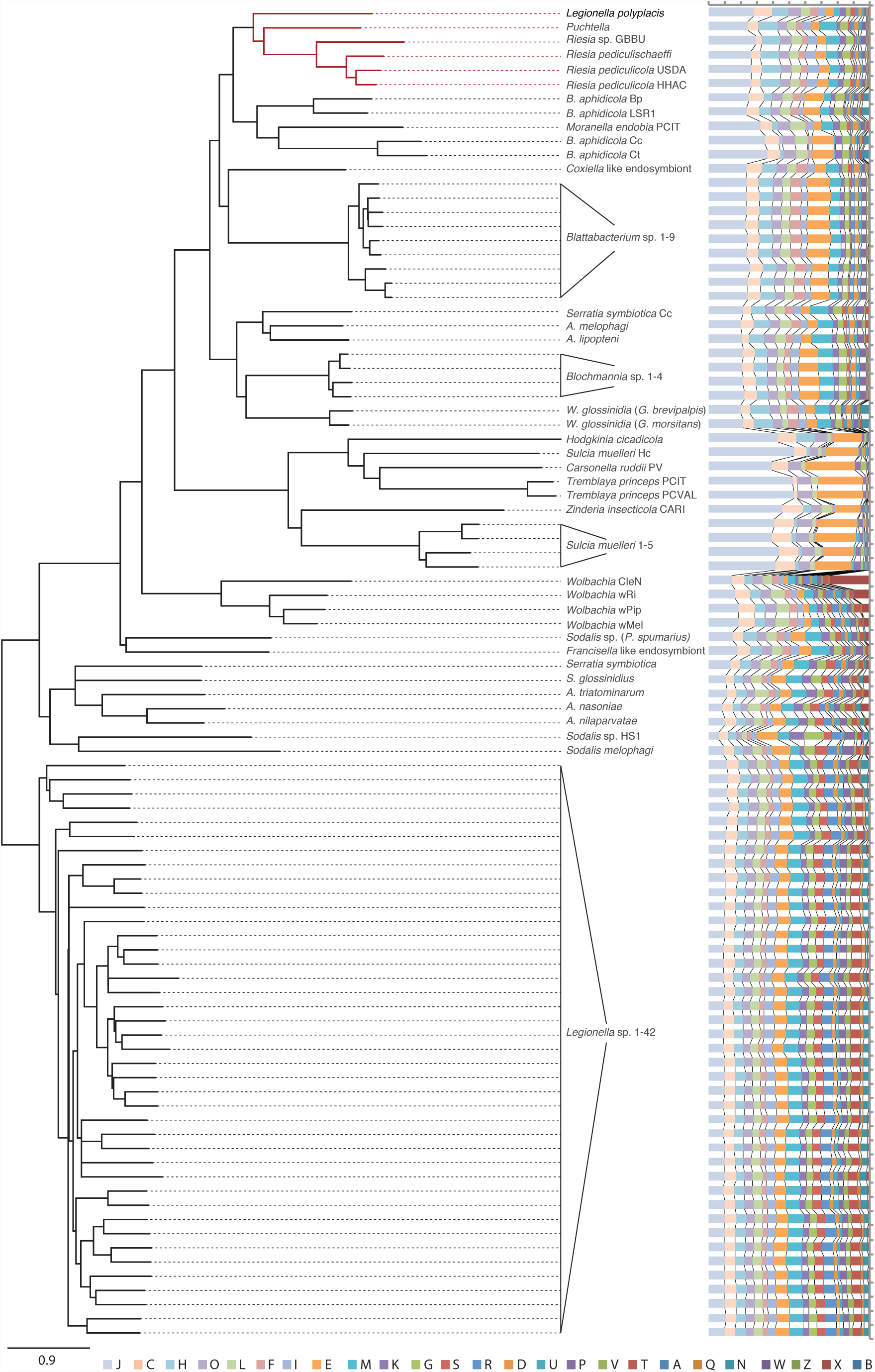
NJ tree calculated for Bray-Curtis distance matrix based on COG orthologs. The bar chart represents COG category content for individual genomes. The categories abbreviations are as follows: **A:** RNA processing and modification, **B**: Chromatin Structure and dynamics, **C:** Energy production and conversion, **D:** Cell cycle control and mitosis, **E:** Amino Acid metabolism and transport, **F:** Nucleotide metabolism and transport, **G:** Carbohydrate metabolism and transport, **H:** Coenzyme metabolism, **I:** Lipid metabolism, **J:** Translation, **K:** Transcription, **L:** Replication and repair, **M:** Cell wall/membrane/envelop biogenesis, **N:** Cell motility, **O:** Post-translational modification, protein turnover, chaperone functions, **P:** Inorganic ion transport and metabolism, **Q:** Secondary Structure, **T:** Signal Transduction, **U:** Intracellular trafficking and secretion, **Y:** Nuclear structure, **Z:** Cytoskeleton, **R:** General Functional Prediction only, **S:** Function Unknown.

**Table 1.** Comparison of the main genomic characteristics^¥^.

|  | *Legionella* spp. | *Legionella polyplacis* | Endosymbionts of hominid lice* |
| --- | --- | --- | --- |
| **Genome size (bp)** | 2,367,087 - 4,818,052 | 529 746 | 528,700 - 580,415 |
| **Number of proteins** | 1,984 - 3,986 | 484 | 444 - 529 |
| **CG content (%)** | 36.60 - 51.10 | 23 | 23.75 - 31.80 |
| **Number of COGs** | 1578 - 2603 | 452 | 428 - 508 |
| **Ø coding density (%)** | 85.43 - 93.10** | 86.50 | 79.17 - 85.96 |
| **Ø gene size (bp)** | 744 - 1,113 | 931 | 849 - 896 |

<sup>‡</sup> The source data are provided in the supplementary file 1 Table S1.
\* The *Riesia phthiripubis* genome was not included into the comparison due to the presence of 444 pseudogenes in the NCBI annotation (CP012846.1).
\*\*Coding densities were calculated only for the species with complete annotations available.

### Metabolism, adaptive processes and HGT

To some degree, however, the convergent evolution/adaptation described above, is limited by phylogenetic constraints, i.e. availability of different metabolic machineries inherited from unrelated bacterial ancestors. For example, both *L. polyplacis* and the *Riesia* species lost capacity to synthetize thiamin. In *Riesia*, this incapacity is compensated by specific thiamin ABC transporter inherited from its ancestor (as inferred from the presence of ABC thiamin transporter in other *Arenophonus* spp.). Boyd, et al. (2017) hypothesized that the loss of thiamin synthesis by *Riesia* in hominid lice and its retention by *Puchtella* in colobus monkey lice may reflect diet differences of the two mammal hosts. They suggest that the complex diet of the hominids makes thiamin available for scavenging by *Riesia*. According to this view, *L. polyplacis* from the rodent associated lice could also scavenge thiamin from its host. However, the specific thiamin transporter is not present in the known *Legionella* genomes and *L. polyplacis* thus lacks both the synthetic pathway as well as specific transporter. Since using the CoFactor database (Fischer, et al. 2010), we determined thiamine as the essential compound required as a cofacor by at least two of the *L. polyplacis* enzymes (transketolase and pyruvate dehydrogenase), it is likely that *L. polyplacis* utilizes some alternative system for its acquisition. As it is known that the ABC transporters can bound to more ligands, we suggest that among possible candidates for this role are the other transporters which could be adapted for nonspecific transfer. For example, the ability to bind thiamin is known for the ABC putrescine transporter (ter Beek, et al. 2014). This transporter is present in *L. polyplacis* and was obviously inherited from its legionellae ancestor.

Further important difference between *L. polyplacis* and *R. pediculicola* introduces horizontal gene transfer (HGT) as yet another determinant of a symbiont’s evolutionary pathway. The significance of HGT for adaptation to a particular life style in bacteria is well recognized. For *Legionella*, Burstein et al. (2016) suggested that many of effectors, with possibly important functions in virulence and intracellular lifestyle, have been recently acquired by this mechanism. Within the system of thousands of predicted effectors, they were able identify only seven effectors shared by all tested *Legionella* species. This, together with the variability of main genome characteristics, shows *Legionella* as highly dynamic system capable of rapid adaptations. Unlike the “free living” legionellae, the HGT in *L. polyplacis* is bound to its symbiotic function known from other blood feeding insects, i.e. provisioning the host with vitamins. This role in *L. polyplacis* is fulfilled by a horizontally acquired complete biotin operon (Fig 2, Supplementary file 3), previously demonstrated for few other bacteria. Since legionellae generally possess the biotin synthesis capacity, the acquisition of the biotin operon by *L. polyplacis* appears as a replacement rather than a de novo acquisition of this function. However, in contrast to this compact six-gene biotin operon, in the genomes of free living legionellae BioC is separated from the rest of the genes. As a remnant of this arrangement, *L. polyplacis* retains the original BioC (position 269231-269893) apart from the complete biotin operon (222799-228219). Various circumstances could possibly drive *L. polyplacis* towards adaptive acceptance of this operon. For example an arrangement of biotin synthesis within a single operon could prove within the highly economized *L. polyplacis* genome more efficient than the genes scattered around the genome as in other legionellae. Also, the acquisition of this operon could follow preceding loss of the biotin synthesis in *L. polyplacis* parasitic ancestor. The best example of such ecological transition due to the HGT acquisition of the biotin operon is a shift of usually parasitic *Wolbachia* to an obligate mutualist in the bed bug *Cimex lectularius* (Nikoh et al. 2014). Altogether, the patterns discussed above show how the combination of different evolutionary forces (phylogenetic constraint, adaptive pressure and HGT) resulted in emergence of a unique genome/phenotype in the genus *Legionella*.

## Acknowledgments

We would like to thank Jana Martinu for providing the lice specimens. This work was supported by the Grant Agency of the Czech Republic (grant 14-07004S to VH). Access to computing and storage facilities owned by parties and projects contributing to the National Grid Infrastructure MetaCentrum provided under the programme “Projects of Large Research, Development, and Innovations Infrastructures” (CESNET LM2015042), is greatly appreciated.

---

**Supplementary file 1:** Tables S1-S3 (Source data for Table 1, List of the compared genomes, *L. polyplacis* genome annotation).

**Supplementary file 2:** Phylogenetic tree inferred by Phyml analysis of concatenated 64-gene matrix. The interrupted branches were shortened by 50%.

**Supplementary file 3: Figures S1-S13.** Phylogenetic trees reconstructed by Phyml (S1-S6) and MrBayes (S7-S12) for individual genes, and by MrBayes for the concatenated matrix (S13; for Phyml analysis of the concatenated matrix see Figure 2).

